# Challenges and Progress in RNA Velocity: Comparative Analysis Across Multiple Biological Contexts

**DOI:** 10.1101/2024.06.25.600667

**Authors:** Sarah Ancheta, Leah Dorman, Guillaume Le Treut, Abel Gurung, Loïc A. Royer, Alejandro Granados, Merlin Lange

## Abstract

Single-cell RNA sequencing is revolutionizing our understanding of cell state dynamics, allowing researchers to observe the progression of individual cells’ transcriptomic profiles over time. Among the computational techniques used to predict future cellular states, RNA velocity has emerged as a predominant tool for modeling transcriptional dynamics. RNA velocity leverages the mRNA maturation process to generate velocity vectors that predict the likely future state of a cell, offering insights into cellular differentiation, aging, and disease progression. Although this technique has shown promise across biological fields, the performance accuracy varies depending on the RNA velocity method and dataset. We established a comparative pipeline and analyzed the performance of five RNA velocity methods on three datasets based on local consistency, method agreement, identification of driver genes, and robustness to sequencing depth. This benchmark provides a resource for scientists to understand the strengths and limitations of different RNA velocity methods.

## Introduction

Single-cell RNA sequencing (scRNA-seq) has enabled the characterization of thousands of transcriptomic states, and many computational methods have been developed to infer the lineages between states. While some cell populations exist in equilibrium, others constantly change due to cell differentiation, environmental changes, cell cycle, or disease perturbations^1^ . During cellular transitions, scRNA-seq data provides a unique opportunity to investigate how cells transition between states and which regulatory programs are responsible for orchestrating such trajectories^2–4^.

Many computational methods exist to infer cellular trajectories from single-cell data, and their performance often varies depending on the type of data, the biological context, and the performance metrics used^5^. One widely used technique, RNA velocity, predicts the future state of a cell based on its mRNA splicing dynamics (Fig. 1a). RNA velocity has been applied to address fundamental questions of cell state transitions in developmental biology^1^ and during perturbation^6–10^. Although RNA velocity has been widely adopted by the community, there are a variety of methods dependent on many parameters and they are known to yield inconsistent or incorrect directionalities^11,12^. Given these limitations, this paper aims to guide researchers in evaluating and choosing the best RNA velocity method for their data.

**Figure 1.**
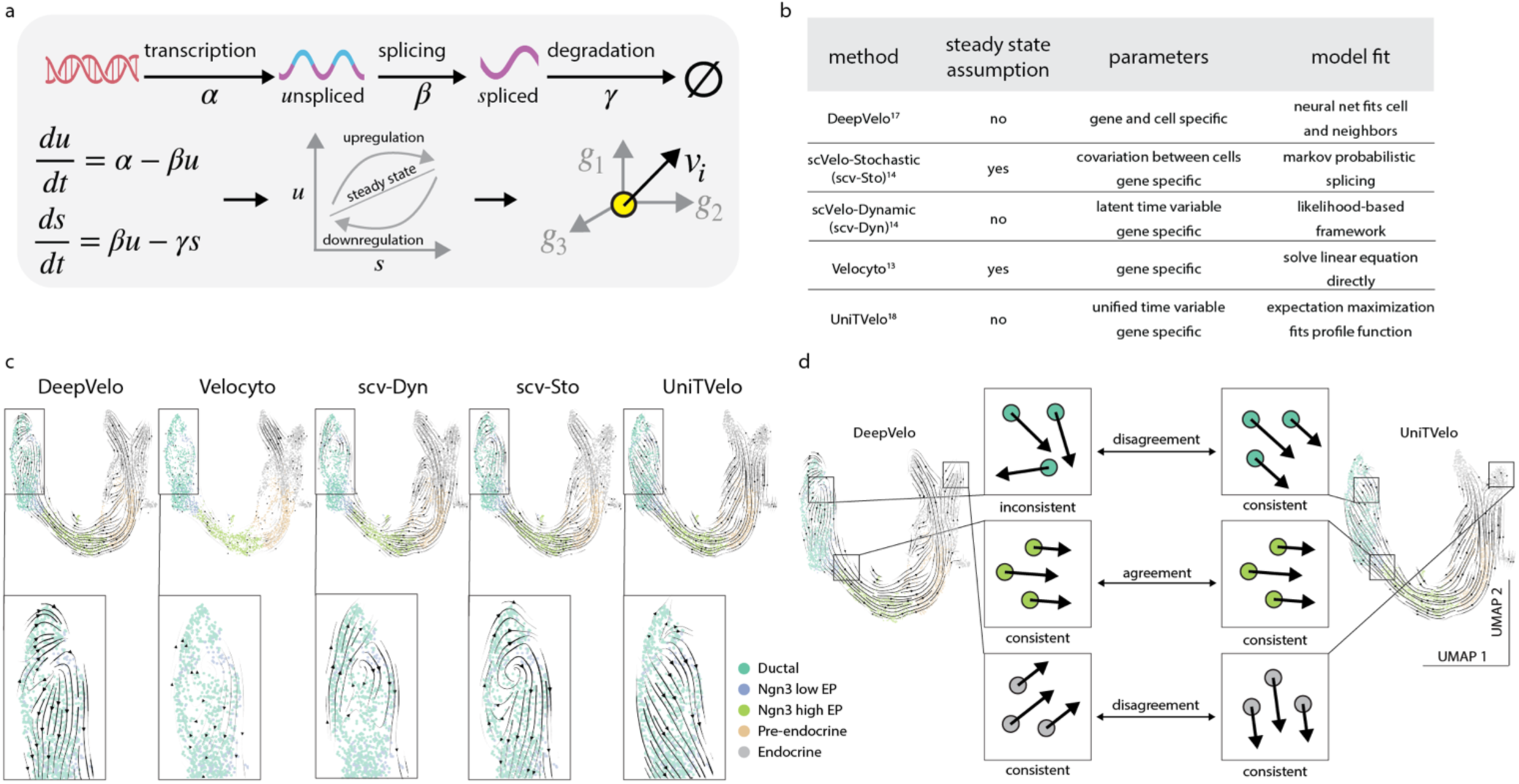
Different RNA velocity methods vary in directionality predictions. a. Overview of the RNA velocity workflow, with the original Velocyto^13^ model as an example. The rates of transcription, splicing and degradation are notated as 𝛼, 𝛽, and 𝛾 respectively. All RNA velocity models use spliced (s) and unspliced (u) mRNA counts as model inputs and predict the directionality of a cell in transcriptomic space. b. Summary of the five RNA velocity models studied in this work and the methodologies implemented in each model. c. UMAP embeddings with RNA velocity predictions of mouse pancreas (n=3696 cells) across the five different methods, highlighting different directionality predictions in the ‘Ductal’ cell type (bottom panel). d. A schematic describing examples of inconsistency and disagreement in the velocity predictions between different methods: (1) the consistency within the single cells in a neighborhood (the colored dots represent single cells), and (2) the agreement in directionality between methods (black arrows between boxes). *See also Figure S1*.

RNA velocity applies dynamic modeling to scRNA-seq data to predict state transitions between individual cells^13,14^. As the mRNA matures in a cell, a fraction of recently synthesized mRNA molecules exist in their unspliced state, while the rest are processed into their spliced, mature state (Fig. 1a, upper panel)^15^. By considering the ratio of spliced and unspliced mRNA measurements, the RNA velocity technique fits a dynamic model to predict the rate of change in the number of mRNA molecules for a specific gene^13^. The rate of change of all genes defines a gradient in the high-dimensional transcriptomic space and predicts the directionality of the molecular states (Fig. 1a, lower panel)^14,15^. The method yields a set of RNA velocity vectors representing the landscape of predicted transitions in the transcriptomic space.

The original RNA velocity model, Velocyto^13^, assumed a constant rate of transcription for each gene and solved a system of linear differential equations with a constant slope for the steady-state solution (Fig. 1b). The model assigns an RNA velocity estimate to each cell based on its deviance from the equilibrium defined by the fit to the linear model. Finally, the directionality in the cell–cell graph, as seen in the UMAP embedding (Fig. 1c), is determined by the similarity of a cell’s future transcriptomic state to other cells in gene space^13^ (Supp. Fig. 1a-c). While Velocyto provided a proof-of-principle and initial approximation to understanding the gene expression landscape, its key assumptions do not always hold^11,12,16^. Not all genes follow the expected behavior of a steady-state model^16^, and the results depend heavily on the chosen hyperparameters and pre-processing decisions^11^.

Recent models have introduced improvements to address these limitations. For example, scVelo introduced a stochastic version (scv-Sto) of the original steady-state model^14^ incorporating stochasticity for transcription, splicing, and degradation, treating them as probabilistic events and resulting in a Markov process (Fig. 1b). Additionally, scVelo proposed a dynamic model (scv-Dyn) to address many of the original issues. scv-Dyn introduces a shared latent time across all cells and fits the gene-specific transcription, splicing, and degradation parameters using likelihood-based expectation maximization^14^ (Fig. 1b).

Like svc-Dyn, UniTVelo defines a shared latent time as a cell-specific component to minimize the discrepancy between the directionality of different genes^17^ (Fig. 1b). Instead of fitting gene-specific splicing functions, UniTVelo^17^ implements a general “profile” function that fits all genes simultaneously, deriving gene-specific splicing parameters in a single step, using expectation maximization. Alternatively, DeepVelo uses a graph convolution network to estimate splicing and degradation rates that are gene and cell-specific^18^. Notably, DeepVelo considers not only a single cell but also its neighbors when fitting the model (Fig. 1b).

Evaluating the accuracy of RNA velocity methods is challenging because ground truth trajectories are rarely available^16^. Moreover, the increasing number of computational methods available make it difficult for scientists to decide the correct workflow for their research (Fig. 1b). For example, even in the mouse pancreas, a well-studied lineage, we observed significant discrepancies in the RNA velocity streams generated by different methods (Fig. 1c inset, e.g., Ductal cells, a progenitor cell state)^19,20^. Therefore, a general benchmark that compares RNA velocity methods in different contexts is necessary to understand this technology’s predictive potential and help scientists choose the best tool to address their questions. We present a comparison of the five RNA velocity methods detailed above (summarized in Fig. 1b). These methods are evaluated across three developmental scRNAseq datasets, including a mouse pancreatic development dataset^14,20^, a single time-point zebrafish 24 hours post-fertilization whole-embryo dataset^21^, and a multi-time point zebrafish neuro-mesodermal progenitors (NMP) lineage dataset^21^ (Supp. Fig. 1d-g).

To analyze the RNA velocity methods, we first evaluate the local consistency (L_c_) within each method, determining if the velocity vectors are consistent across neighbor cells with high transcriptomic similarity (Fig. 1d). Second, we examine method agreement (A_1_ and A_2_) to assess the landscape’s robustness across different RNA velocity methods (Fig. 1d). Disparate or contradictory results from various RNA velocity methods undermine our confidence in the predicted trajectories. We extend this framework and evaluate the concordance in the downstream identification of driver genes. Finally, we analyze the robustness of each method relative to the number of reads, simulating the sensitivity of RNA velocity methods to sequencing depth. We observe that the smoothness and robustness of RNA velocity landscapes vary significantly depending on the cell type and biological context. We expect our benchmark to provide insight into the strengths and weaknesses of RNA velocity as a tool for understanding cell fate dynamics during differentiation.

## Results

### Local Consistency

We initially evaluated the methods by analyzing the consistency of velocity vectors within neighborhoods, a common metric for evaluating the performance of RNA velocity methods^17,18,22,23^. The molecular state transitions taking place can be represented with a Markov transition matrix between individual cells that considers the directionality and strength predicted by RNA velocity^14,15,24^. In the Markov transition matrix, each cell has a state transition vector representing the transition probabilities that determine the cell’s likely future state. For each cell, we quantified the local consistency as the relative alignment between a cell’s state transition vector and those of the most transcriptionally similar cells. We defined the metric L_C_ as the average cosine similarity between a cell and its 30 nearest neighbors (Fig. 2a)^18^. Under this definition, neighbor cells with transition vectors in inconsistent directions will have a low score, whereas agreement between neighbors will result in higher consistency scores (Fig. 1d).

**Figure 2.**
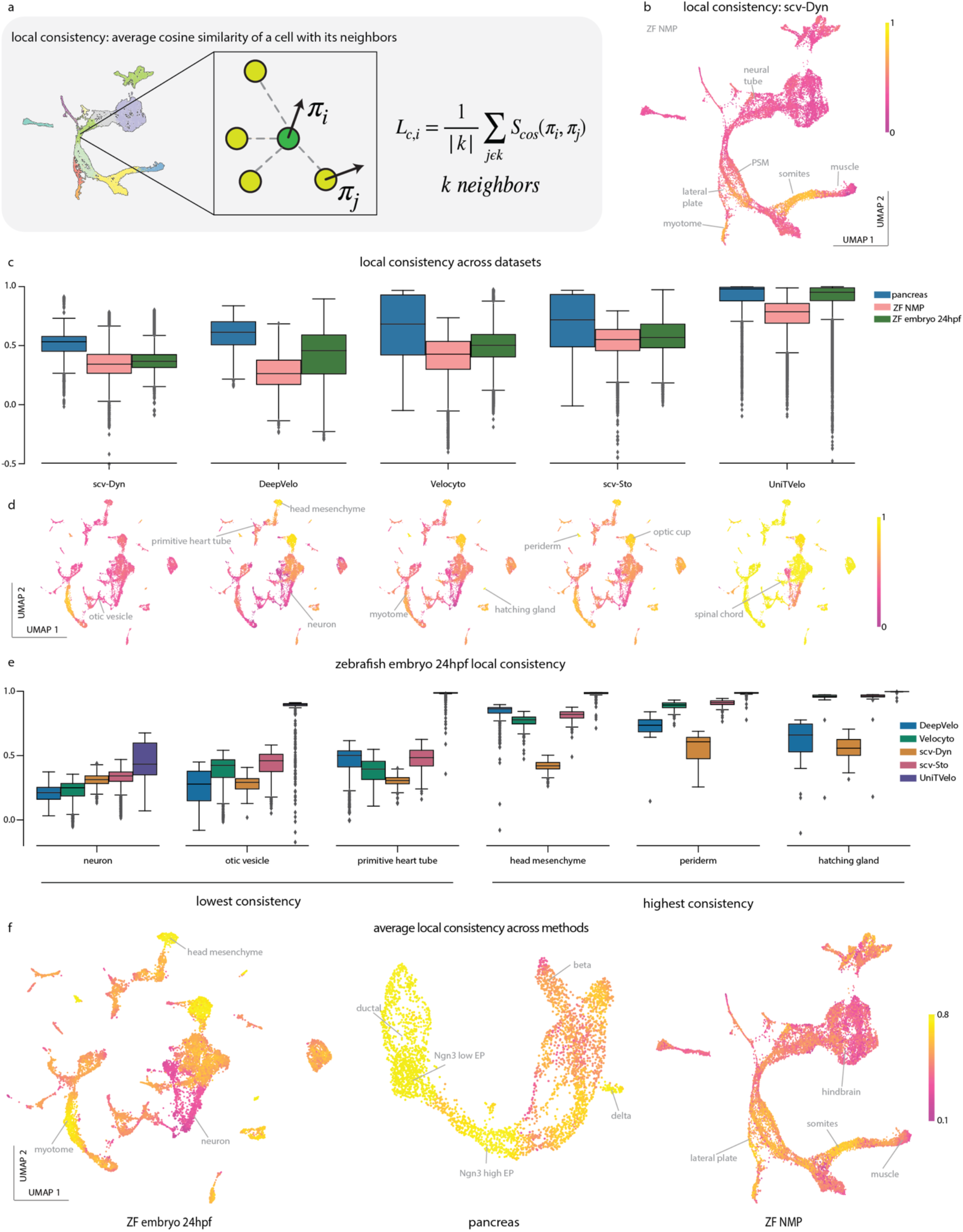
RNA velocity local consistency in cell neighborhoods. a. The local consistency quantifies the average velocity transition vector alignment of a cell with its nearest neighbors. Each dot is a cell, and the local consistency is the average of the cosine similarities between the target cell and each of its nearest neighbors (k=30). b. Single-cell local consistency of scv-Dyn projected into the UMAP embedding for the ZF NMP dataset (n=16035 cells). c. Local consistency distributions for each RNA velocity method across the three datasets. d. UMAP embeddings for zebrafish whole-embryos 24hpf (n=12914 cells) colored by the single-cell local consistency calculated for each RNA velocity method. The labels highlight cell populations with high or low consistency. e. Local consistency distributions for the three cell types with the highest and three with the lowest average local consistency in the zebrafish embryos 24hpf dataset. f. UMAPs colored by the average (consensus) local consistency across all methods, for the three datasets. The labels highlight cell populations with high or low consensus consistency. *See also Figure S2*.

We next calculated the L_C_ scores for the cells in the zebrafish neuromesodermal progenitor (ZF NMP) lineage dataset for the RNA velocity method scv-Dyn and projected them on the UMAP embedding (Fig. 2b). The distribution of L_C_ scores showed significant differences across cell types and UMAP regions. Cell types from well-defined lineages, such as the mesodermal-derived cells, showed high consistency, in particular the axial mesoderm (PSM → somites → muscle)^25,26^, and the lateral plate mesoderm (Fig. 2b). In contrast, those with more complex cellular heterogeneity, such as neural cells (see Fig. 2b, neural tube and hindbrain), showed the lowest consistency values. More generally, the L_C_ distribution showed high heterogeneity and appeared to be cell type-specific (Fig. 2b, Supp. Fig. 2).

We then compared the local consistency distributions for all three datasets and across RNA velocity methods (Fig. 2c, Supp. Fig. 2). Some methods, such as UniTVelo, showed high L_C_ for all cells in the dataset, indicating a high degree of smoothness across the whole velocity graph, independently of the cell type (Fig. 2c, Supp. Fig. 2a, c). In contrast, DeepVelo and scv-Sto showed intermediate L_C_ values with heterogeneous distributions for all three datasets (Fig. 2c, d, Supp. Fig. 2a, c). We observed a clear trend in L_C_ across methods and datasets, where scv-Dyn consistently showed lower values, and UniTVelo showed high L_C_ values across all datasets (Fig. 2c).

To understand the heterogeneity in L_C_ values, we explored the distributions for the cell types in each dataset in more detail. In whole zebrafish embryos at 24 hours post fertilization (ZF embryo 24hpf) the distribution of L_C_ showed significant differences between cell types (Fig. 2d), except for UniTVelo’s velocity calculations, which resulted in high L_C_ values across diverse cell types (Fig. 2e, Supp. Fig. 2b, d).

For the cell types with the highest L_C_ values, we observed strong agreement across most methods, suggesting that the RNA velocity signal in these neighborhoods is strong enough to be discernible by different models (Fig. 2e, Supp. 2b, d). Together, these results indicate that the landscape’s smoothness varies depending on cell type.

Given the diverse outcomes from different methods, we grouped the results by creating a consensus score, the average L_C_ across all methods, which enabled the characterization of overall high or low agreement for different datasets (Fig. 2f). In the pancreas dataset, 92% of cells exhibited consensus L_C_ above 0.5, indicating high agreement between methods (Fig. 2f). Similarly, 74% of cells in ZF embryo 24hpf dataset had a consensus L_C_ above 0.5, but only 39% of cells are over 0.5 in the ZF NMP dataset (Fig. 2f). The low consensus in the ZF NMPs dataset can be explained by the heterogeneity in the age of the cells, as this dataset integrates multiple time points. The consensus L_C_ could assist in identifying regions where the differentiation signal is strong enough to reconstruct single-cell trajectories based on RNA velocity. On the other hand, lower L_C_ across all methods could indicate that the velocity signal for the cell type is noisy, their differentiation process is more complex, or the cells are not differentiating (e.g., hindbrain cells in the ZF NMP dataset or neurons in the ZF embryo 24hpf (Fig. 2e, Fig. Supp. 2b). Overall, we observed that well-defined developmental transitions with low cell diversity have high local consistency, whereas lineages with complex diversity show low local consistency.

### Method Agreement

Though local consistency with a method is an important metric in evaluating RNA velocity methods, alternatively, one can ask if the vector predictions from different velocity methods agree. Agreement between methods can help to identify lineages and cellular states with stronger velocity signals that correlate with biological relevance. We analyzed the agreement between methods using two approaches: (1) comparing the directionality of each cell’s vector predictions for each pair of methods and (2) comparing each method to the ‘median vector,’ the central vector derived from all methods (Fig. 3a).

**Figure 3.**
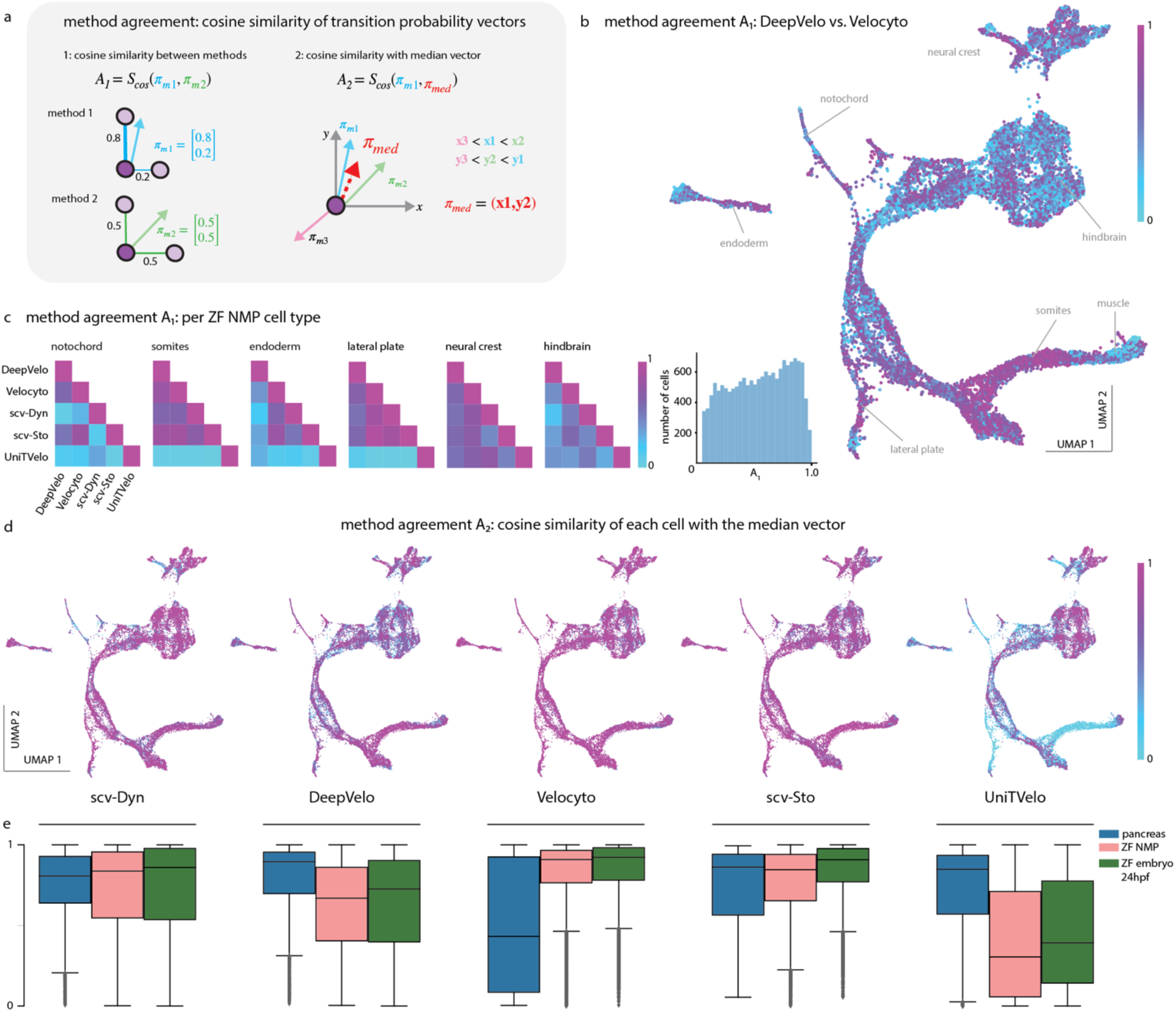
Comparing single-cell velocity agreement between methods. a. Each RNA velocity method yields a cell-cell transition graph. Part (1) of the schematic shows different transition probability vectors for the same cell obtained from different methods. A transition vector is defined by the transition probabilities between cell states in the graph. The method agreement (1) metric quantifies the similarity between cell transition vectors across datasets using cosine similarity. Part (2) of the schematic illustrates the median (central) vector for each cell, computed by taking the median of all transition vectors across methods. The method agreement (2) metric quantifies the similarity of cell transition vectors from each method as compared to the median vector by using cosine similarity. b. UMAP embedding for the ZF NMP dataset with the method agreement (1) between DeepVelo and Velocyto. The labels highlight cell populations with high or low agreement between the two methods. The histogram shows the distribution of method agreement values for all the cells. c. Pairwise comparisons for six cell types from the ZF NMP dataset across all methods. The heatmap shows the mean method agreement across individual cells within a cell type for each pair of methods. d. UMAP embeddings for ZF NMP for each RNA velocity method, colored by each method’s agreement (2) with the median vector. e. Distribution of method agreement (2), each method’s agreement with the median vector for the three datasets. *See also Figures S3, S4, S5, and S6*.

The metric A_1_ is defined as the landscape’s agreement across pairs of different methods (Fig. 3a, right, Supp. Fig. 3-5, 6a). More specifically, the agreement (A_1_) quantifies the cosine similarity between a pair of transition probability vectors obtained from different methods for the same cell. In comparing DeepVelo and Velocyto in the ZF NMP dataset, levels of agreement varied by cell type. We projected the A_1_ distribution on the UMAP embedding and observed high agreement in the mesodermal lineages and lower agreement in the hindbrain cells, consistent with patterns of developmental heterogeneity^21^ (see Results Local Consistency, Fig. 3b). The mesodermal lineage showed high agreement only in the early stages of differentiation, but as cells differentiate into muscle, the agreement across methods dropped to almost zero (Fig. 3b). The A_1_ (DeepVelo vs. Velocyto) scores (Fig. 3b, histogram) over all cells were widely distributed, indicating heterogeneity in the agreement between the two methods, with patterns of low and high agreement similar to those found for local consistency in the ZF NMP dataset (i.e. highest in somites, lowest in the hindbrain) (Fig. 2b).

Next, we investigated the agreement across methods for cell types in the ZF NMP dataset and observed consensus across most methods. To examine the method agreement for different cell types more closely, we computed the mean A_1_ for each cell type across all pairs of methods (Fig. 3c).

Analysis of the agreement between methods by cell type revealed varying levels of agreement depending on the method and cell population (Supp. Fig. 3-5). The lateral plate, somites, and endoderm exhibited concordance among all methods except for UniTVelo (Fig. 3c). Examining the UMAP for the ZF NMP dataset revealed that UniTVelo had predicted the opposite direction to the known biological trajectory, going from the differentiated cell type towards the progenitor^21,27^ (Supp. Fig. 1b). For three cell types (notochord, endoderm and hindbrain) scv-Dyn and DeepVelo had low agreement, which is correlated with cell diversity and transcriptomic complexity. (Fig. 3c, Supp. Fig. 6b).

To expand the investigation of method agreement on a global scale and evaluate the systematic disagreement across methods, we compared each method to the median transition vector of each cell, computed across all methods (Fig. 3a, right). We then computed the cosine similarity of each method’s transition vector with the cell’s derived median vector (A_2_) (Fig. 3a). The A_2_ metric, therefore, provides a measure of how well each method agrees with the consensus prediction across all methods. For the ZF NMP dataset, we noticed that UniTVelo systematically disagreed across many cell types, whereas DeepVelo had a scattering of cells with low A_2_, mixed with higher values (Fig. 3d). Velocyto, scv-Dyn, and scv-Sto had high cosine similarity with the median vector (Supp. Fig. 6c, d).

When applying the analysis to the three datasets, we found method agreement with the median vector varied depending on the dataset, except for scv-Dyn and scv-Sto (Fig. 3e). DeepVelo had high levels of agreement for the pancreas dataset, but lower levels of agreement in the zebrafish datasets (pancreas: median A_2_=0.894, ZF NMP: median A_2_=0.667, ZF embryo 24hpf: median A_2_=0.726) (Fig. 3e). The agreement for UniTVelo followed a similar pattern, with much lower levels of agreement in the zebrafish datasets (pancreas: median A_2_=0.848, ZF NMP: median A_2_=0.305 and ZF embryo 24hpf: median A_2_=0.392) (Fig. 3e). The high performance of deep learning-based methods (DeepVelo and UniTVelo) on the pancreas (a commonly tested dataset for developing RNA velocity methods) can indicate overtraining or over smoothing^17,18^. The opposite pattern was seen in Velocyto, where performance was low on the pancreas dataset (median A_2_=0.432), and the method achieved the highest agreement across all methods on the zebrafish datasets (ZF NMP: median A2=0.908, ZF embryo 24hpf: median A2=0.922) (Fig. 3e). Altogether, the variation in agreement across datasets and methods underscores the importance of implementing and comparing predictions across multiple methods when interpreting RNA velocity.

### Downstream: Overlap of Driver Genes

We next explored how the disagreements between methods are propagated in downstream analysis by evaluating the overlap in macrostates and top driver genes in the pancreas dataset, the most well-studied lineage among the datasets we evaluated. We utilized CellRank to identify macrostates and driver genes, i.e. genes whose expression highly correlates with a specific trajectory or lineage with a velocity kernel generated from each method^15^ (see Methods). CellRank estimates absorption probabilities (i.e. probability of a cell fate trajectory towards a particular terminal state) using ensembles of random walks. Genes are classified as drivers if they are systematically highly expressed in cells that are more likely to differentiate towards a given terminal state^15^ (see Methods).

While the macrostates identified by CellRank generally agreed across methods and corresponded to the Leiden clusters and cell type annotations (initial cluster Ductal, terminal Alpha, Beta, Delta, and Epsilon), we found that some states didn’t agree across methods (Fig. 4a, b, Supp. Fig. 7a, see Delta, Ngn3 low EP, Ductal 5).

**Figure 4.**
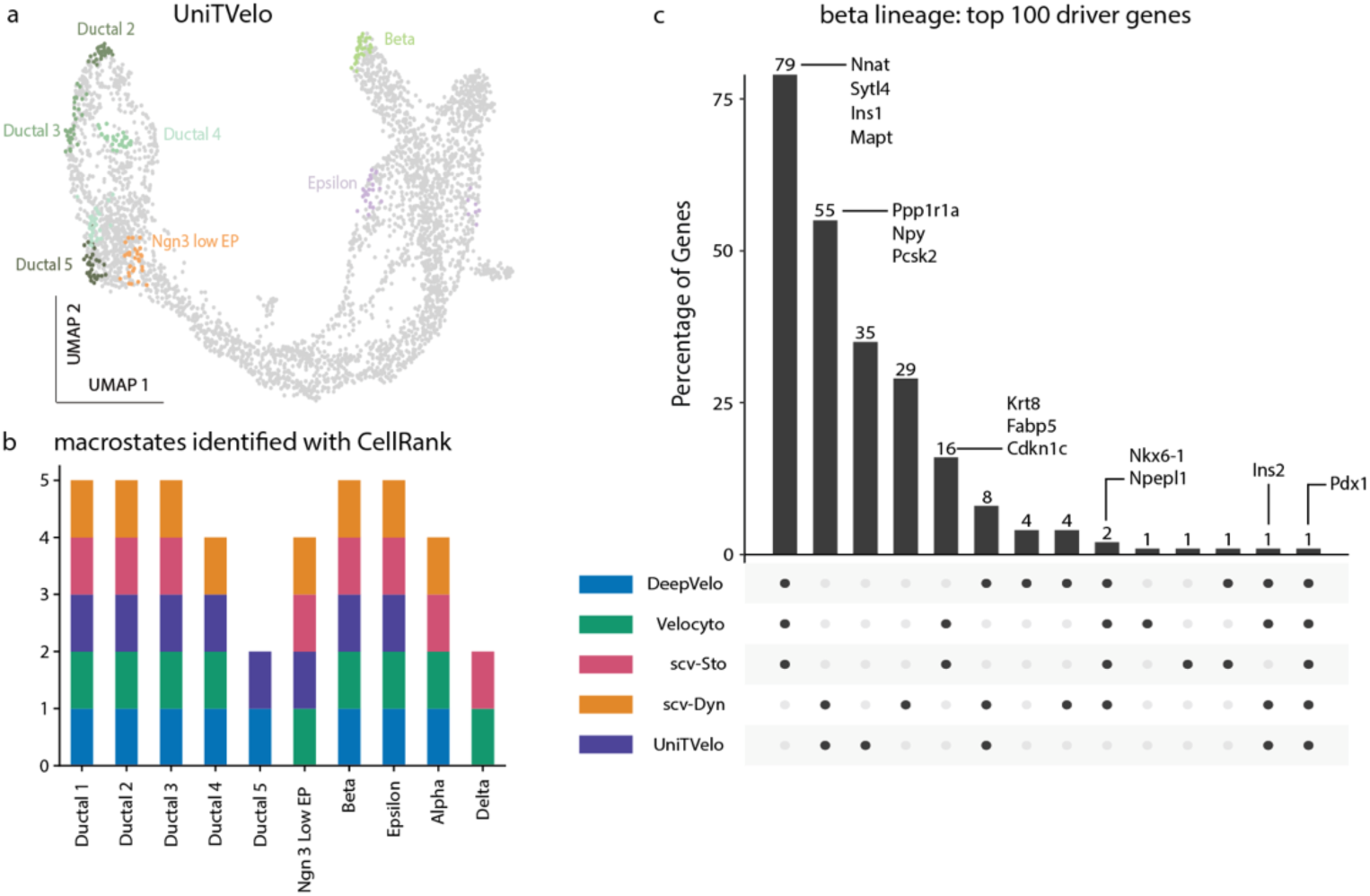
Comparing driver genes predicted from different methods. a. Macrostates identified by CellRank based on the velocity kernel from UniTVelo in the pancreas dataset (see Methods). b. A summary of the macrostates identified by CellRank for different RNA velocity methods. The x-axis shows the macrostate label. The y-axis shows the number of methods in which the macrostate was identified by CellRank. c. A histogram showing the top 100 genes for the Beta lineage from each method. The bars indicate the number of genes that appear at the intersection of the methods. *See also Figure S7*.

In exploring the role of driver genes, we focused on the terminal state Beta, as it was robustly identified by CellRank in all methods (Fig. 4b). We looked at the overlap among the identified top 100 driver genes across the methods. Strikingly, we found only one gene, Pdx1, at the intersection of all methods (Fig. 4c). Pdx1 is an essential transcription factor and master regulator in the development and maintenance of Beta cells^28^. While it is encouraging that all methods agreed on Pdx1, it is notable that only one gene appeared in the intersection.

Interestingly, the different methods clustered into two groups with regard to the driver genes identified (Fig. 4c). The first group (DeepVelo, scv-Sto and Velocyto, agreed on 79% of driver genes) (Fig. 4c); whereas the second group (scv-Dyn and UniTVelo) agreed on 55% of their predicted driver genes (Fig. 4c). scv-Dyn and UniTVelo both utilize a shared latent time variable to fit their models. When conducting an analysis of the driver genes for the Epsilon terminal state, we found similarly low levels of overlap, with three genes at the intersection of all methods (Supp. Fig. 7b). The general lack of agreement in driver genes emphasizes the importance of considering multiple methods when making trajectory predictions with RNA Velocity and testing these predictions using experimental perturbations or other validation methods.

### Robustness to Sequencing Depth

The robustness of the RNA velocity methods to changes in sequencing depth was analyzed to evaluate the sensitivity of the methods. To simulate different levels of depth, we subset the ZF embryo 24hpf dataset, randomly selecting different proportions of reads (2, 5, 12, 25, 50, 80, 95 and 98%, each repeated five times), computed each velocity method and evaluated the robustness of the prediction as compared to the full dataset (Fig. 5a, Supp. Fig. 8-9). The impact on the velocity predictions can be observed at a high level, as trajectories vary with different proportions of reads. When computed by DeepVelo with 2% of the reads, the velocity flow for the myotome is in the opposing direction as compared to all increased subsets (Fig. 5b, Supp. Fig. 8a). With Velocyto, the flow through the hindbrain appears to become more complex as the proportion of reads increases (Fig. 5c, Supp. Fig. 8c).

**Figure 5.**
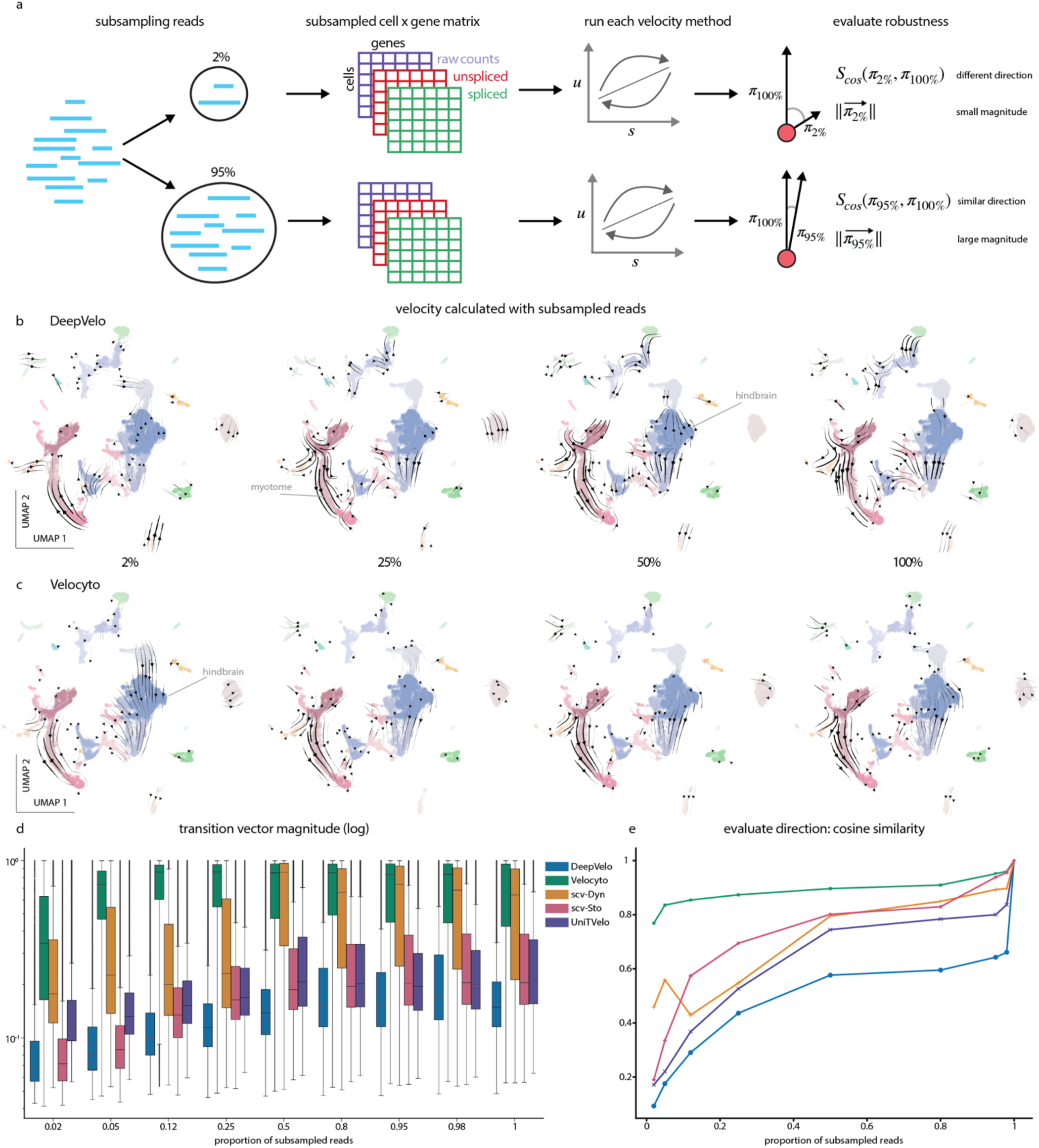
Robustness of methods to sequencing depth. a. Sequencing depth is simulated by taking a randomized subset of the reads, including 2, 5, 12, 25, 50, 80, 95, and 98%. The input matrices, including raw, spliced, and unspliced counts, are then derived from the reads. All RNA velocity methods are run on the subset, and we evaluate the robustness based on the vector’s magnitude and direction as compared with the predictions from 100% of the reads. b. Illustration of UMAP embeddings of the ZF whole-embryo 24hpf with RNA velocity predictions from DeepVelo, calculated from subsets 2, 25, 50, and 100% of the reads. The labels highlight cell populations with directionality disagreement between subsets. c. UMAP embeddings of the ZF whole-embryo 24hpf with RNA velocity predictions from Velocyto, calculated from subsets 2, 25, 50, and 100% of the reads. The labels highlight cell populations with directionality disagreement between subsets. d. Distribution of transition vector magnitudes across all subsets for the five RNA velocity methods for the ZF whole-embryo 24hpf dataset. e. Comparison of directionality robustness for each method, determined by calculating the cosine similarity between the transition vector from the subset and the directionality of the transition vector derived from 100% of the reads. This calculation was averaged across all cells in the ZF whole-embryo 24hpf dataset *See also Figures S8, S9 and S10*.

To evaluate the sensitivity of each method to sequencing depth, we examined the magnitude and direction of the transition vectors. As expected across all methods, the magnitude of the transition vectors generally increased with the number of reads (Fig. 5d). The transition vectors from Velocyto had the largest magnitudes across all levels, and scv-Dyn had similarly high magnitudes starting at 50% of the original sequencing depth (Fig. 5d). The transition vector magnitudes for scv-Sto, UniTVelo, and DeepVelo remained much smaller (Fig. 5d).

In comparing the magnitudes derived from subsets to those calculated from 100% of the data, the vector magnitudes converged for scv-Sto and Velocyto, whereas DeepVelo, scv-Dyn, and UniTVelo maintained low levels of correlation with the magnitudes from the full reads (Supp. Fig. 10a). Together, the data indicates that more reads lead to larger transition vectors and that the vector magnitudes for Velocyto and scv-Sto are more robust to read numbers (Supp. Fig. 10a).

We evaluated the directionality robustness of the transition vectors by computing the cosine similarity between each cell’s predicted transition vector from the subset reads and the predicted vector from the full reads. Velocyto was the most robust, with the largest increase at 5% of the reads to a cosine similarity around 0.85, after which additional reads provided a small amount of improvement (Fig. 5e). All other methods plateau at lower values, with a jump at the end between 98% and 100% of the reads and reaching their stable points around 50% of the reads (Fig. 5e). DeepVelo was the lowest, reaching a plateau of ∼0.5, while scv-Sto and scv-Dyn were just above UniTVelo (at 0.7) (Fig. 5e). We checked that the magnitude did not affect the evaluation of directionality and observed that the variance was smallest for Velocyto (Supp. Fig. 10b, c). The directionality of the velocity predictions from Velocyto are the most robust to simulated lower sequencing depth.

## Discussion

We evaluated the performance of five RNA velocity methods on three developmental datasets by analyzing their local consistency, method agreement, overlap of driver genes, and robustness to sequencing depth. Collectively, the RNA velocity methods identified known biological trajectories and important driver genes, with each method displaying varying levels of performance depending on the dataset and evaluation metric. Our research emphasizes the importance of implementing a method that best fits the dataset and encourages the utilization of multiple approaches when identifying trajectories for further experimentation.

Local consistency varied across cell types and methods (Fig 2c, 2e). For cell types with high local consistency, all five methods showed similar results, suggesting that the signal in the data was high enough to be identified by different models. In some terminal state cell types (i.e., muscle, hindbrain, neural tube), we observed low consistency, which may be due to noisy measurements or heterogeneity in subpopulations^16^ (summarized in Fig. 6). Many methods benchmark their performance using metrics to measure local consistency^18,22,23^ and our findings highlight the value of incorporating different analytical perspectives.

**Figure 6.**
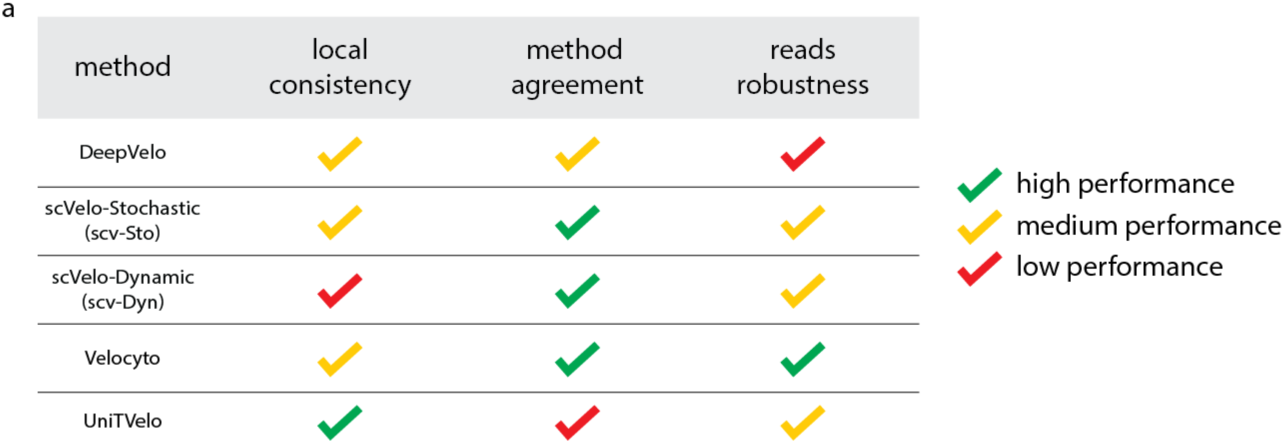
Guidelines to choose an RNA Velocity Method. Table of metrics used to evaluate each RNA velocity method, with color indicating high (green), medium (yellow) or low (red) performance of the method for each metric. Local consistency indicated is the median local consistency for each method across the median for each dataset: high (0.8-1), medium (0.5-0.8), and low (0-0.5). Method agreement is the median agreement with the center vector for each method, across the median for each dataset: high (0.8-1), medium (0.5-0.8), low (0-0.5). Reads robustness is based on the cosine similarity plateau value for the directionality of each method: high (0.8-1), medium (0.5-0.8), low (0-0.5).

When evaluating the methods in comparison to the median vector as computed across all methods, the inconsistencies for different datasets were apparent. Because the pancreas dataset is often used as a benchmark dataset for RNA velocity methods, the high performance of other methods over Velocyto (the original method) could indicate improvement in the field^18,29^. UniTVelo had low method agreement for both zebrafish datasets yet high local consistency, which is likely due to over-smoothing as the method fits a single profile function for all cells and genes and utilizes a unified latent time to infer dynamics in a sample^17^ (Fig. 6). Both deep learning-based methods, UniTVelo and DeepVelo, had lower method agreement in the zebrafish datasets as compared to the pancreas (Fig. 3e). We suspect that their default parameters were optimized for a specific training set, and that with parameter optimization these methods may perform more accurately, as deep learning models are more complex^30^. The downstream analysis identification of driver genes was sensitive to the differences in velocity calculations between methods, emphasizing the need to include multiple RNA velocity predictions before making decisions about further experimentation.

The levels of robustness to the sequencing depth, as simulated by downsampling the number of reads, varied depending on the model type of each method. The deep learning-based approaches were more sensitive to the number of reads, as the models require a larger amount of input data^30^. Velocyto was the most robust to the number of reads (Fig. 6). For scientists who would like to implement RNA velocity, we recommend considering sequencing depth when considering the best method for their data.

Several limitations apply to the findings reported here: 1) the number of datasets is limited and only includes two organisms with vastly different numbers of cells, 2) our study is not comprehensive of all current RNA velocity methods, and 3) all comparisons rely on default parameters as suggested by the authors as we do not explore method optimization. Nonetheless, we believe our framework provides an initial approach to address these questions.

With the current restraints of scRNA-seq, short-read mRNA sequencing alone might be insufficient to describe the differentiation dynamics of these cell types and additional “omic” modalities could improve the modeling of velocities^16^. For example, recent approaches combine RNA sequencing with ATAC-seq to jointly model RNA velocity^31^. Other single-cell techniques may lead to improvements in RNA velocity models, such as long-read sequencing^32,33^. Capturing the full splicing dynamics with long-read sequencing, along with improvements in genome annotation, may provide additional transcriptomic information needed to more accurately predict the future state of a cell with RNA velocity. While RNA velocity is a predictive model and ground truth is unavailable, comparing multiple methods can provide overall insights into trajectories in the transcriptomic space to help us understand the underlying biological processes.

## Acknowledgments

We thank the Data Science and Royer teams of the Chan Zuckerberg Biohub for their helpful discussion. We are grateful for the review and feedback from Sandy Schmid, Jordão Bragantini, Joan Wong and Yang-Joon Kim. The Chan Zuckerberg Initiative through the Chan Zuckerberg Biohub San Francisco (CZB SF) provided funding for this work.

## Methods

### Code availability

All relevant code is available here https://github.com/czbiohub-sf/comparison-RNAVelo.

### Single-cell quality control and RNA velocity method implementation

The mouse pancreatic developmental dataset is available through scVelo^14,20^. The ZF NMP and the ZF embryo 24hpf datasets are subsets of the Zebrahub data^21^. For each RNA velocity method, we followed the workflow and preprocessing as outlined in each method’s tutorial. We executed the same preprocessing for Velocyto, scv-Sto, scv-Dyn, and DeepVelo. For each dataset, we normalized the raw counts using scVelo v0.2.5. Genes with fewer than 20 total detected counts were excluded. Gene counts were normalized by dividing by the total counts per cell and multiplying by the median total counts per cell. The top 2,000 highly variable genes were then log normalized using the functions scvelo.pp_filter_genes, scvelo.pp.normalize_per_cell, scvelo.pp.filter_genes_dispersion, and scvelo.pp.log1p respectively. Utilizing the scVelo pipeline, we computed first and second moments for the velocity estimation, with 30 principal components and 30 nearest neighbors (sc.pp.neighbors, scvelo.pp.moments). The scVelo v0.2.5 Deterministic mode recapitulates the Velocyto steady-state model. We ran this model with scvelo.tl.velocity mode=’deterministic’ using default parameters. For scv-Sto and scv-Dyn, we ran scvelo.tl.velocity (scVelo v0.2.5) with mode=’stochastic’ and mode=‘dynamic’ with default parameters. We ran DeepVelo^18^ (0.2.5rc1) with the default configuration. For UniTVelo^17^ (v0.2.5.2), all preprocessing is part of the model, and we ran the model using its default parameters. We input precomputed Leiden clusters for the input parameter ‘cluster,’ as calculated with scanpy for the zebrafish datasets^21^. For the pancreas dataset, we input the cell type parameter, as given with the dataset, as the ‘cluster’ input.

### Velocity counts and generation of the velocity graph

For each of the RNA velocity methods, a ‘velocity’ prediction was generated. This yields a matrix in which each cell has a velocity vector with directionality in the transcriptomic space, and a ‘velocity_graph,’ a cell state–cell state graph. To compare the methods directly, we used the scVelo v0.2.4 function scv.utils.get_transition_matrix to compute cell state–cell state transition probabilities from the velocity graph, based on the similarity of a cell’s predicted future state to the profile of cells observed in the sample. We input the dataset and each method’s ‘velocity_graph’ variable to get the transition matrix.

### Consistency within single-cell neighborhoods

For each method, we calculated the local neighborhood consistency (L_C_) as shown in DeepVelo^18^. We calculate 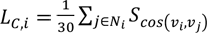 where 𝐿*_Ci_* indicates the local consistency of the cell i, and 𝑁_#_ indicates the 30 nearest neighbors of the cell. We computed the cosine similarity between the cell state transition vectors of each cell to its neighbors, using np.inner and np.linalg.norm from numpy v1.23.5. L_C_ is defined as the average cosine similarity across all 30 nearest neighbors. We used the local neighborhood calculation for each RNA velocity method, and for all datasets. To identify cell types with higher or lower consistency, we grouped cells together by Leiden cluster, and labeled each Leiden cluster with the most prominent cell type represented. To evaluate the local neighborhood consistency across all methods, we took the average L_C_ across all RNA velocity methods for each single cell.

### Transition vector agreement between methods

We calculated the pairwise method agreement by computing the cosine similarity between two transition vectors for the cell from transition matrices created by two different RNA velocity methods: *A*_1_ = *Scos*(𝑣*_i,M_*_1_, 𝑣*_i,M_*_2_),, where *Scos(*𝑣*_i,M_*_1_*, _vi,M_*_2_) indicates the cosine similarity between state transitions vectors for cell i, with pairs of methods M1 and M2 for each cell. The cosine similarity is defined as the norm of each vector, which we implemented using np.inner and np.linalg.norm with numpy v1.23.5. For each pair of methods and each cell, this yields a method agreement score A_1_.

With the transition matrix from each method, we calculate the central vector for each cell as the median transition vector across all methods. We then compute the agreement with the central vector for each individual method as the cosine similarity 𝐴_1_ = 𝑆*_cos_* (𝑣*_i, M1_,* 𝑣*_i, Med_*) of the transition vector 𝑣*_i,M1_* (for the method M1 and cell i), with the median vector for the cell i, 𝑣*_i,Med_*.

### Computing macrostates and driver genes with CellRank

In the pancreas dataset, we used the variable ‘velocity,’ the velocity for each gene and cell generated from each RNA velocity method, to create the velocity kernel from CellRank^15^ v2.0.0, and we then computed a transition matrix with CellRank’s default parameters, including model=’deterministic’, similarity=’correlation’, and softmax_scale=6.925. The velocity kernel was generated with the parameters backward=False and vkey=’velocity.’ Next, we followed the CellRank pipeline and generated macrostates, defined the terminal states, and computed lineage drivers. We fit the pancreas cluster_key=’clusters’ as the cell type variable, and we set the number of macrostates to 8.

Based on the macrostates generated, we set the terminal states to pancreatic cell types ‘Beta’, ‘Alpha’, ‘Delta’, and ‘Epsilon’ if identified. We then computed the lineage drivers for the Beta lineage, as the terminal state Beta was found in all methods.

In CellRank, the lineage driver genes were identified by the correlation of gene expression with the fate probabilities of the cells from the lineage. To find the overlapping driver genes between velocity methods, we selected the top 100 driver genes with the highest correlation to the Beta lineage from each method, and we utilized UpSet^34^ to visualize the overlap between all groupings of methods.

### Sampling reads to simulate robustness to sequencing depth

To simulate robustness, we subsampled the reads, starting from the bam files for the zebrafish embryo 24 hours post fertilization. The full dataset contains four zebrafish embryos. For each embryo, we utilized samtools to subset a percentage (2, 12, 25, 50, 80, 95, and 98%) of the total reads, creating 5 randomly sampled replicates for each percent. We then ran Velocyto^13^ v. 0.17.17 on each subsampled bam file for each embryo, with the reference genome used for the original dataset^21^. We combined the resulting count matrices for the four fish at each subsampling percentage and iteration and ran the five RNA velocity methods on each of the resulting 35 anndata objects as outlined earlier in the methods.

We analyzed the directionality and robustness by comparing the resulting transition matrices, utilizing the same workflow outlined in the method section ‘velocity counts and generation of the velocity graph’ to generate the matrix. We then computed the magnitude of the transition vector for each cell (np.linalg.norm), and the velocity vector (generated by RNA velocity in transcriptomic space). To analyze the directionality, we computed the cosine similarity for each cell between its transition vector generated by a proportion of the reads and the transition vector generated by 100% of the reads, repeated for each replicate. The results were averaged across cells and replicates before plotting.

To ensure the evaluation of directionality was not largely affected by vectors with a small magnitude, we also computed a min/max weighting for each cosine similarity value, multiplying by the minimum magnitude of the two vectors and dividing by the maximum magnitude of the vectors.

---

**Supplementary Figure 1.**
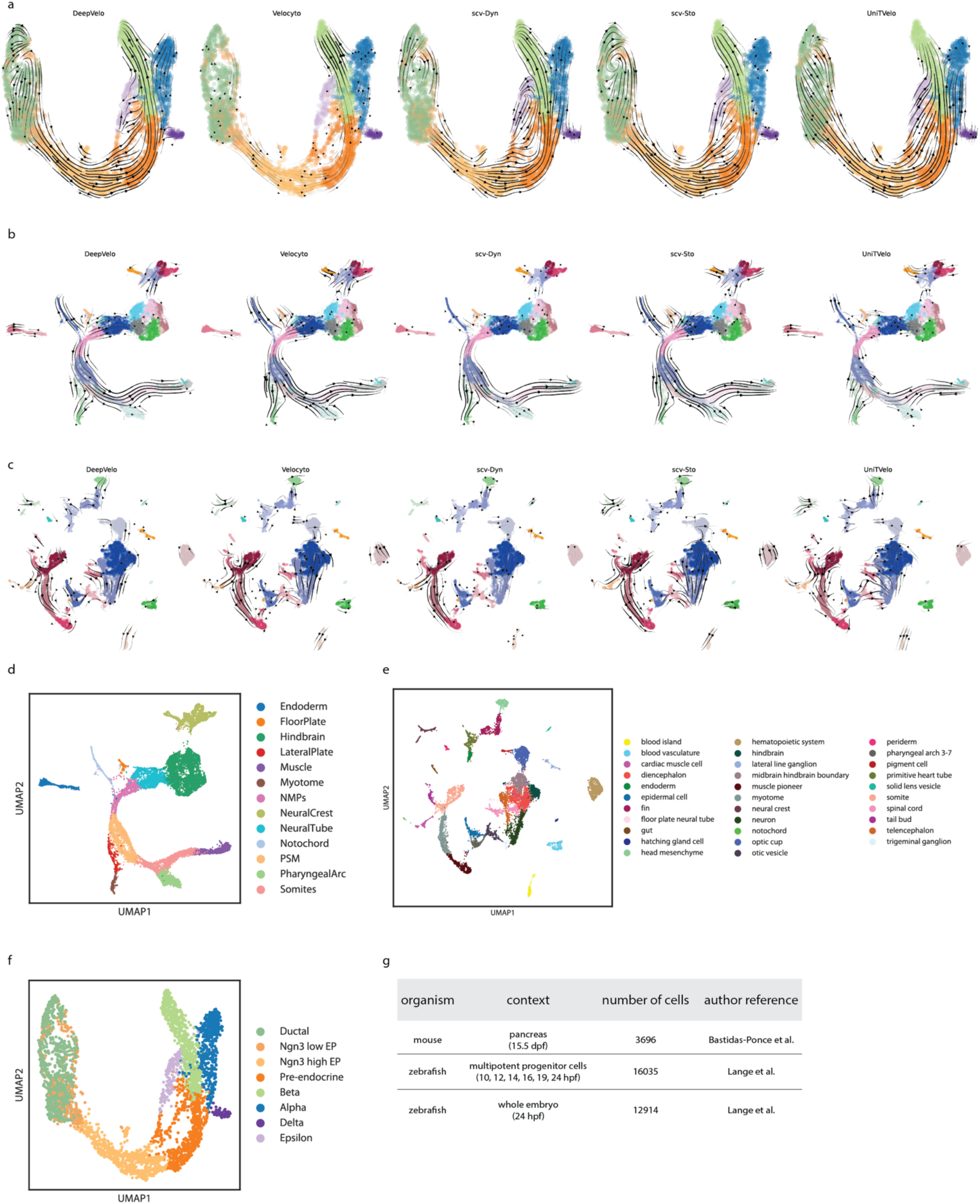
RNA Velocity methods and datasets, associated with Figure 1. a. RNA velocity UMAP projections for five methods, implemented in the pancreas dataset b. RNA velocity UMAP projections for five methods, implemented in the zebrafish NMP (ZF NMP) dataset c. RNA velocity UMAP projections for five methods, implemented in the zebrafish full embryo 24 hours post fertilization (ZF embryo 24hpf) dataset d. ZF NMP UMAP colored by cell type annotations e. ZF embryo 24hpf UMAP colored by cell type annotations f. Pancreas UMAP colored by cell type annotations g. Table of datasets used in the paper, including information about the organism, biological context, number of cells, and author reference.

**Supplementary Figure 2.**
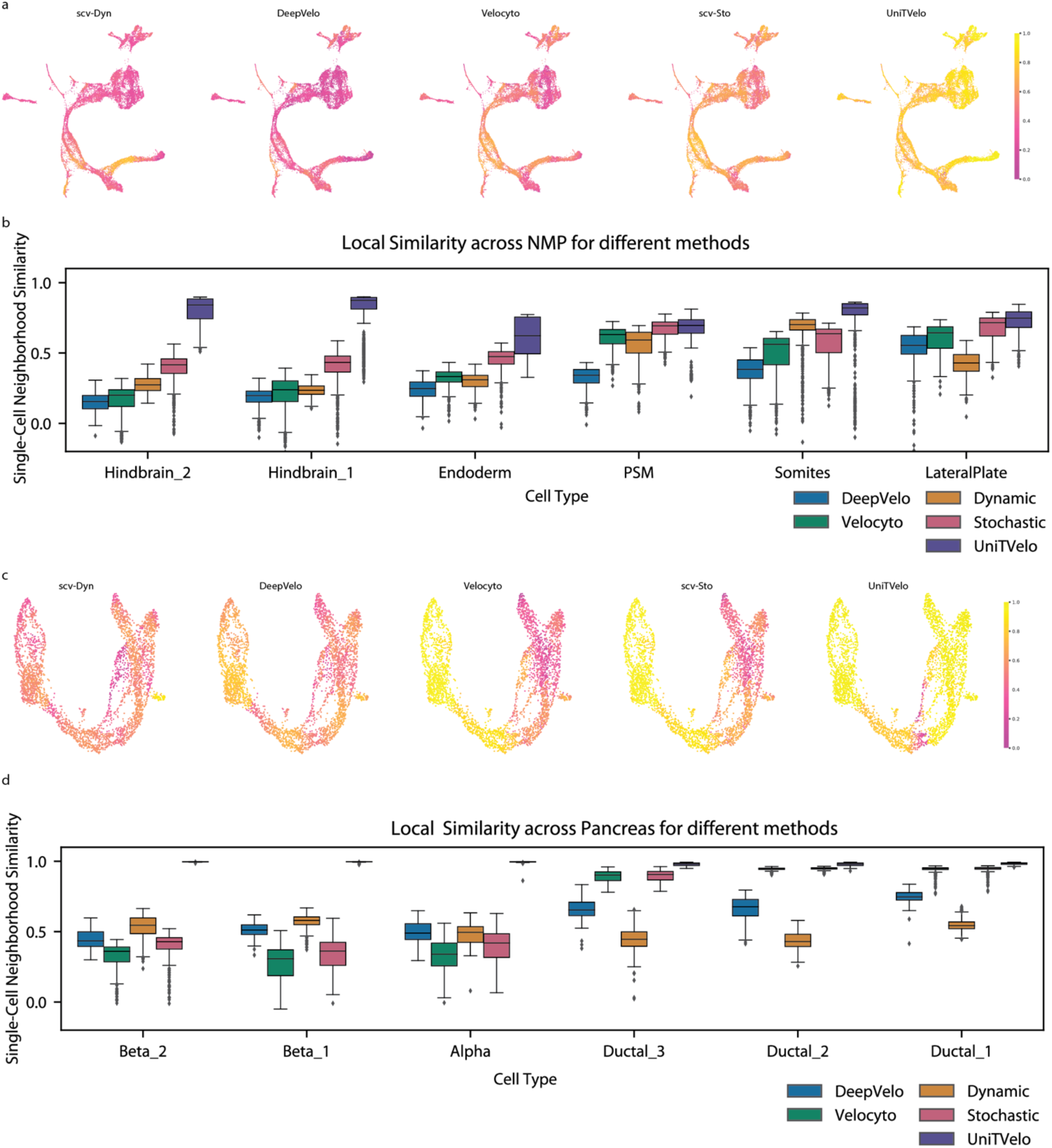
Local consistency across the ZF NMP and pancreas datasets, associated with Figure 2. a. UMAP embeddings for ZF NMP dataset colored by the single-cell local consistency for each RNA velocity method. b. Local consistency distributions for the three top and bottom cell types from the ZF NMP dataset, as ranked by average local consistency. c. UMAP embeddings for pancreas dataset colored by the single-cell local consistency for each RNA velocity method. d. Local consistency distributions for the three top and bottom cell types from the pancreas dataset, as ranked by average local consistency.

**Supplementary Figure 3.**
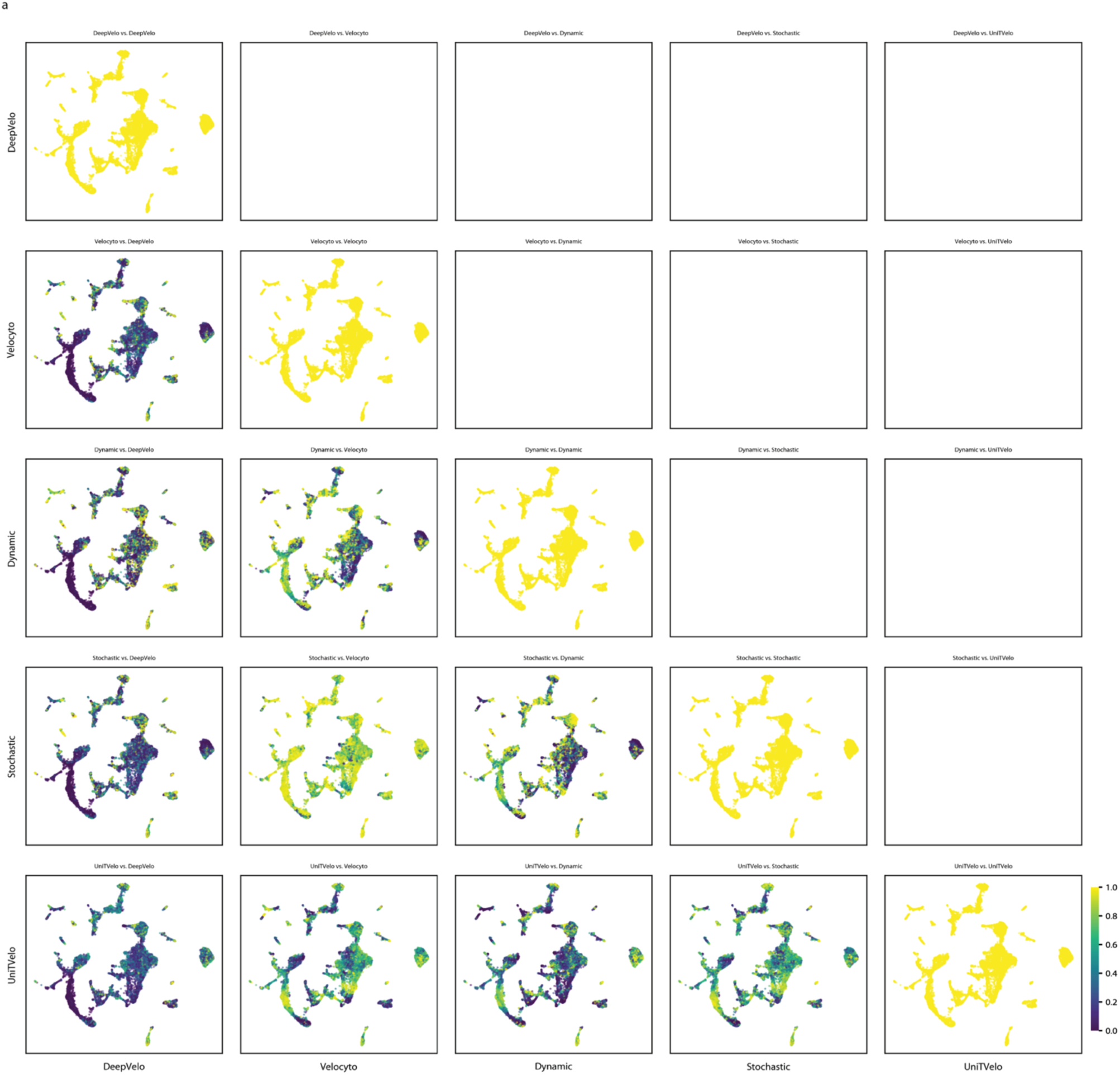
Pairwise Method Agreement UMAPs for ZF embryo 24hpf, associated with Figure 3 UMAP embedding for the ZF embryo 24hpf dataset with all unique pairs of method agreement.

**Supplementary Figure 4.**
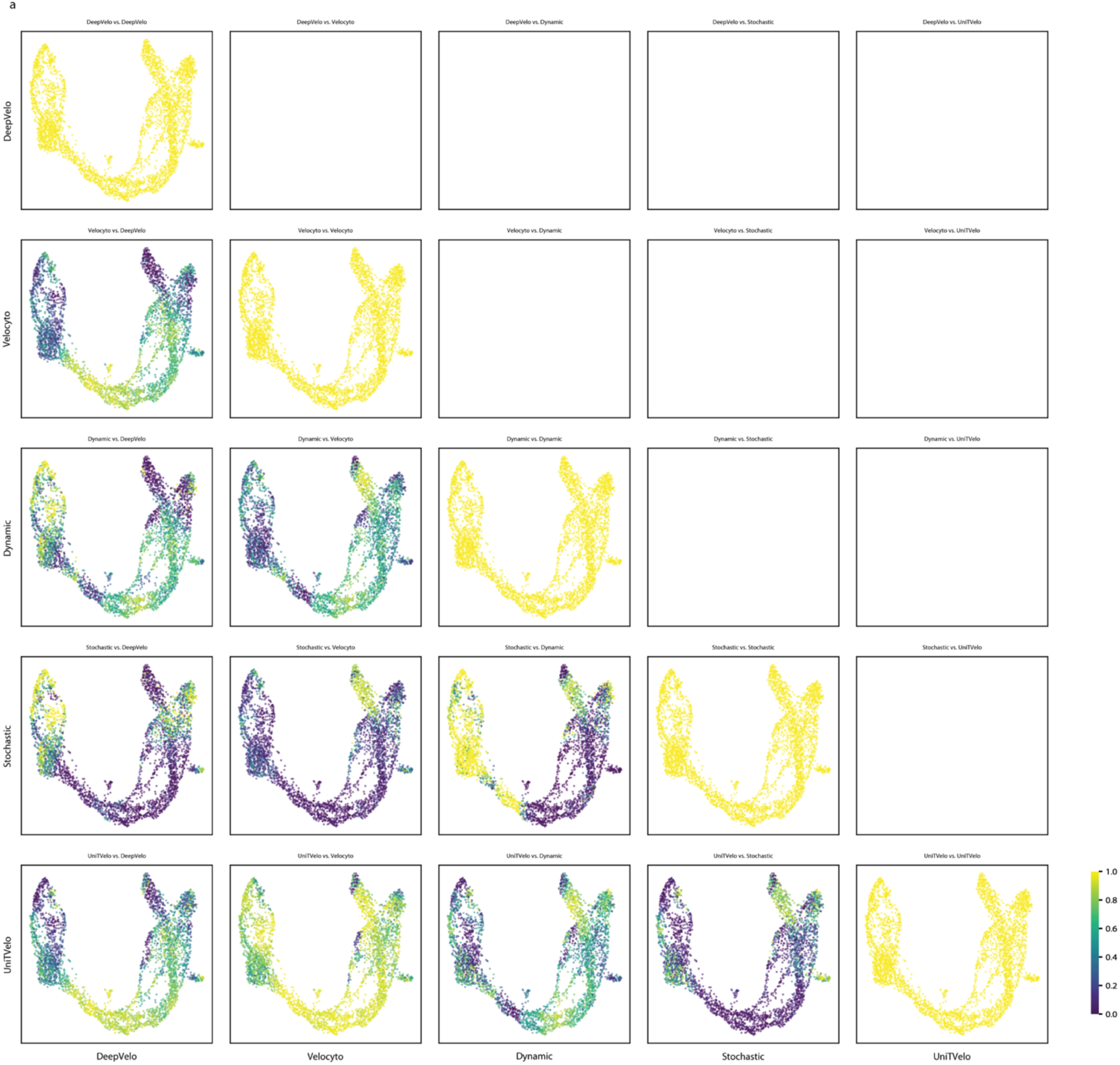
Pairwise Method Agreement UMAPs for pancreas, associated with Figure 3 UMAP embedding for the pancreas dataset with all unique pairs of method agreement.

**Supplementary Figure 5.**
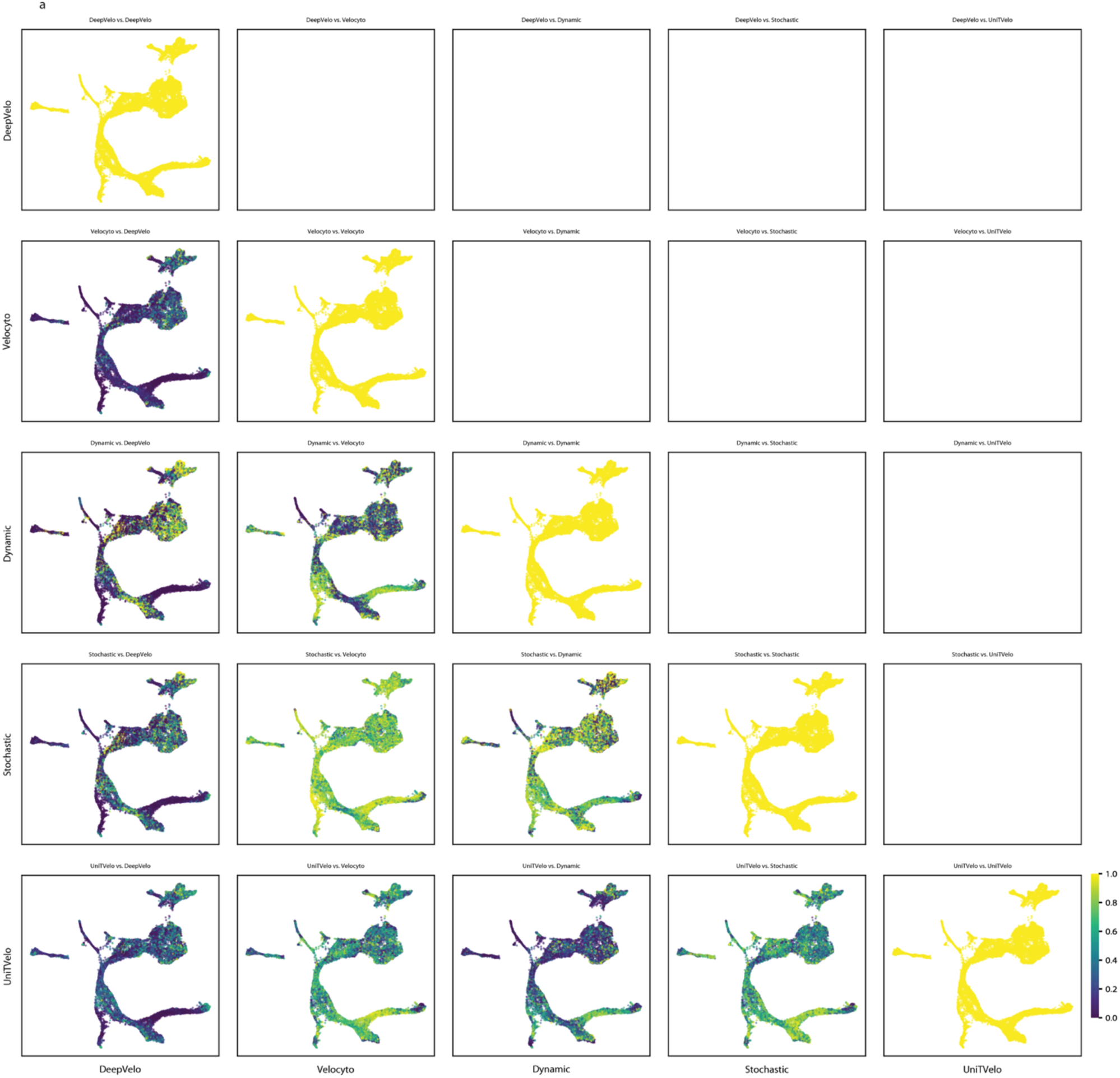
Pairwise Method Agreement UMAPs for ZF NMP, associated with Figure 3 UMAP embedding for the ZF NMP dataset with all unique pairs of method agreement.

**Supplementary Figure 6.**
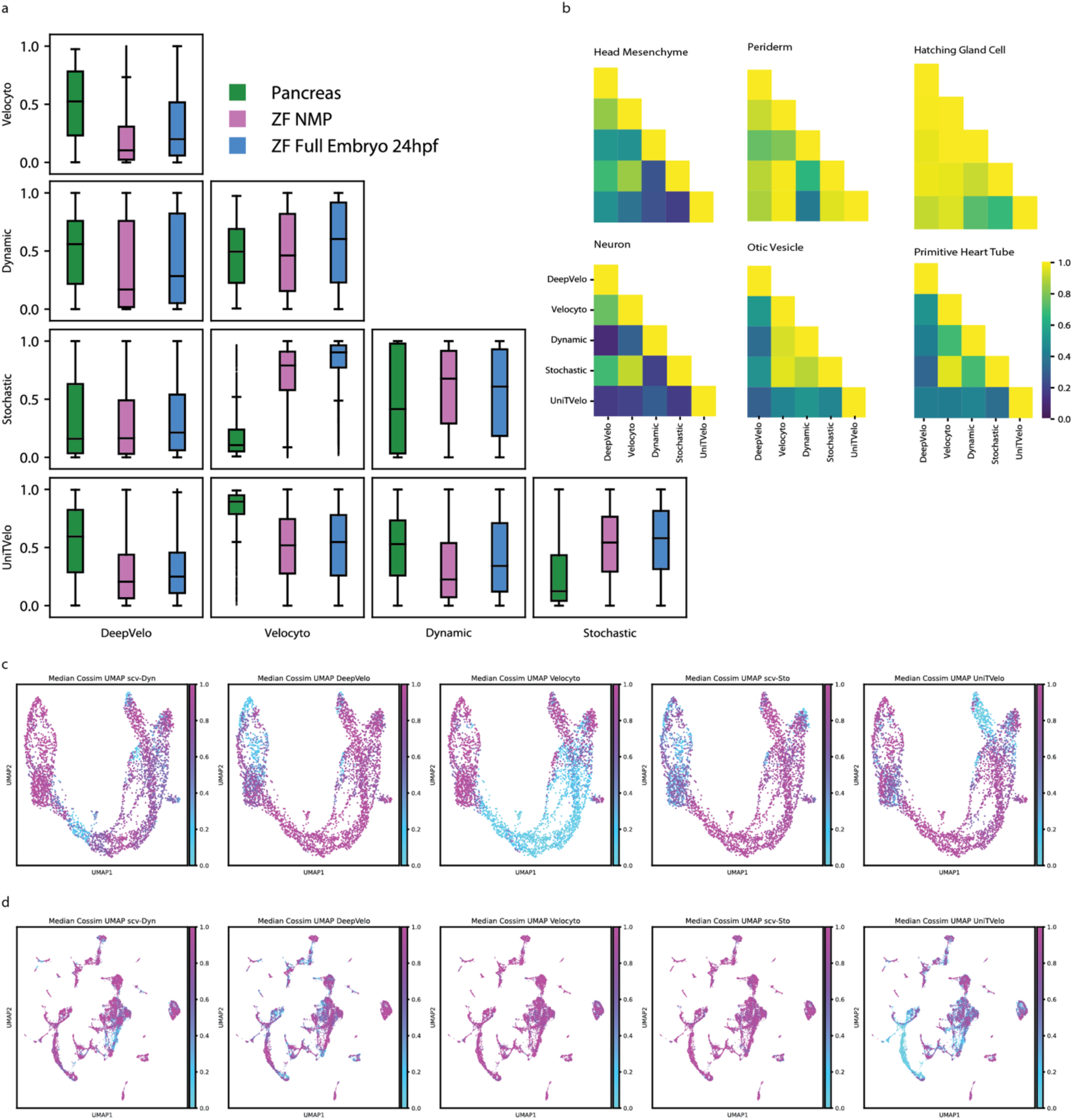
Additional pairwise method agreement distributions and agreement with median vector, associated with Figure 3. a. Boxplots with distributions of pairwise method agreement for each pair of method for each of the three datasets. b. Pairwise comparisons for six cell types from the ZF embryo 24hpf dataset across all methods. The heatmap shows the median method agreement across individual cells within a cell type for each pair of methods. c. UMAP embeddings for the pancreas for each RNA velocity method, colored by each method’s agreement (2) with the median vector. d. UMAP embeddings for Zebrafish NMPs for each RNA velocity method, colored by each method’s agreement (2) with the median vector.

**Supplementary Figure 7.**
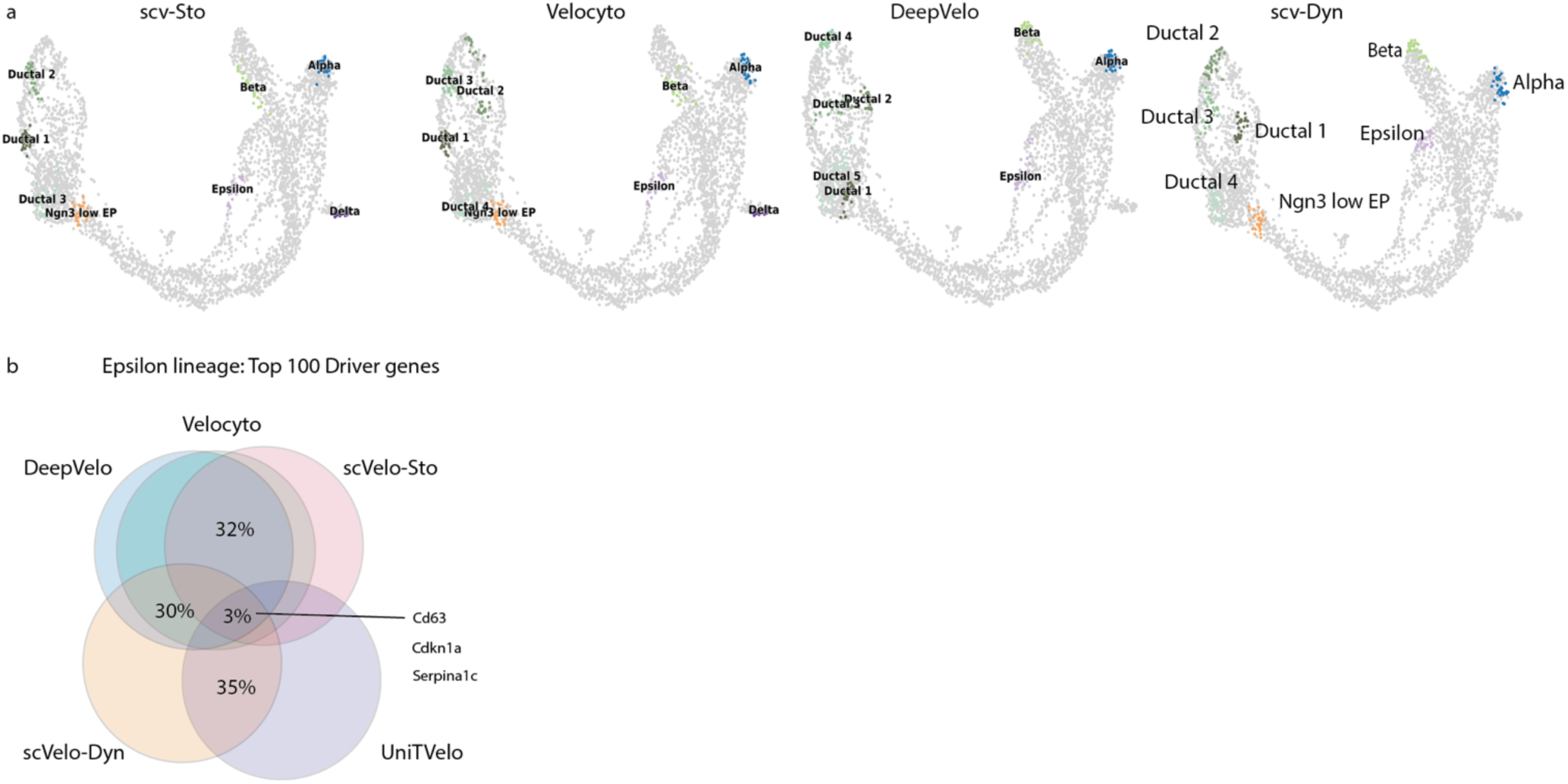
Overlap of Driver Genes and CellRank Terminal States, associated with Figure 4. a. Macrostates identified by CellRank for scv-Sto, Velocyto, DeepVelo and scv-Dyn in the pancreas dataset. CellRank identified different macrostates for each dataset depending on the model’s predictions (see Methods). b. Venn diagram with the percentage overlap across all methods and select groups indicated of the top 100 driver genes for the Epsilon lineage. Genes identified across all methods as top driver genes are labeled.

**Supplementary Figure 8.**
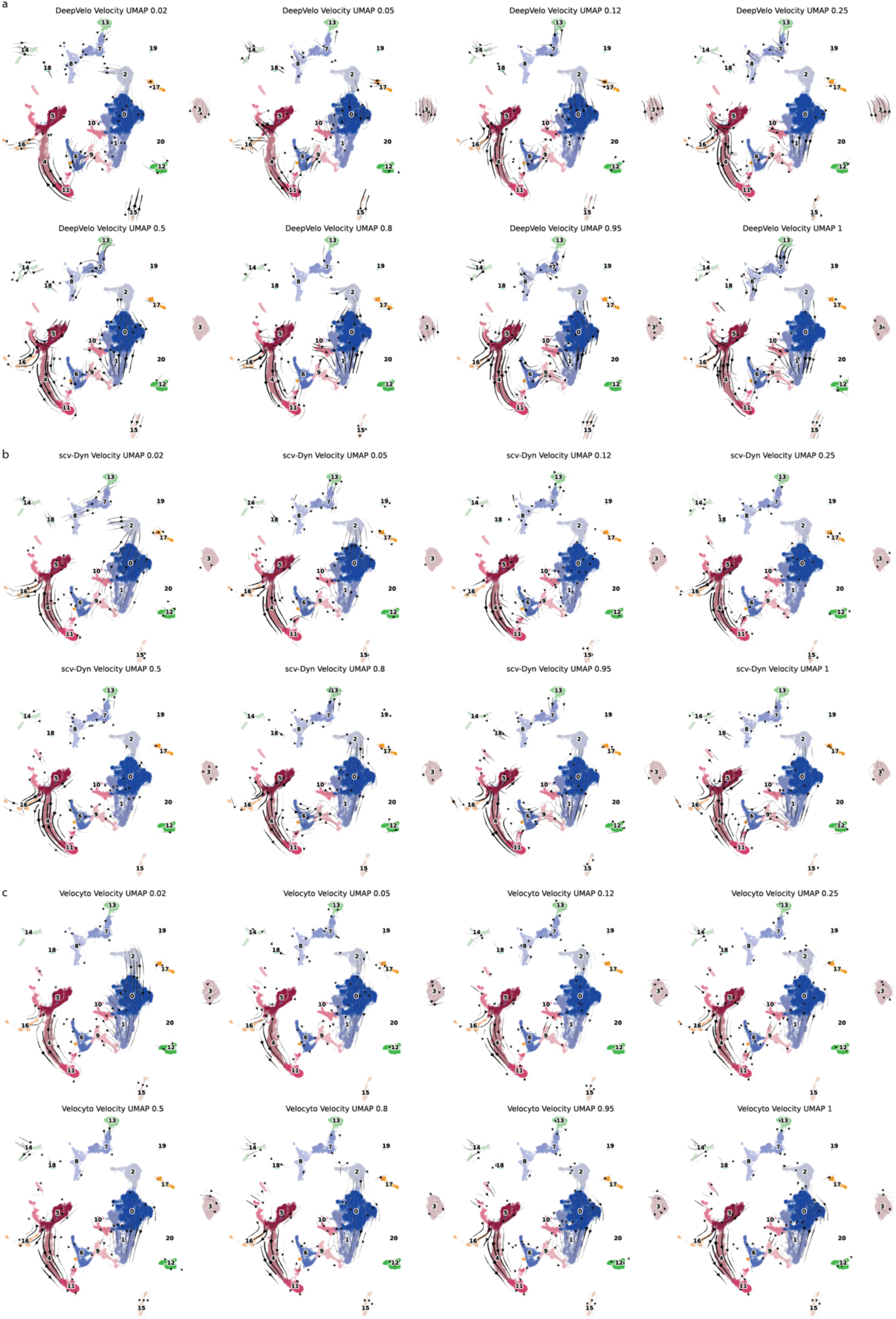
Velocity UMAPs with Subsets of Reads for DeepVelo, scv-Dyn and Velocyto, associated with Figure 5. a. UMAP embeddings of the ZF 24hpf whole-embryo with RNA velocity predictions from DeepVelo, calculated from subsets 2, 5, 12, 25, 50, 80, 95 and 100% of the reads. b. UMAP embeddings of the ZF 24hpf whole-embryo with RNA velocity predictions from scv-Dyn, calculated from subsets 2, 5, 12, 25, 50, 80, 95 and 100% of the reads. c. UMAP embeddings of the ZF 24hpf whole-embryo with RNA velocity predictions from Velocyto, calculated from subsets 2, 5, 12, 25, 50, 80, 95 and 100% of the reads.

**Supplementary Figure 9.**
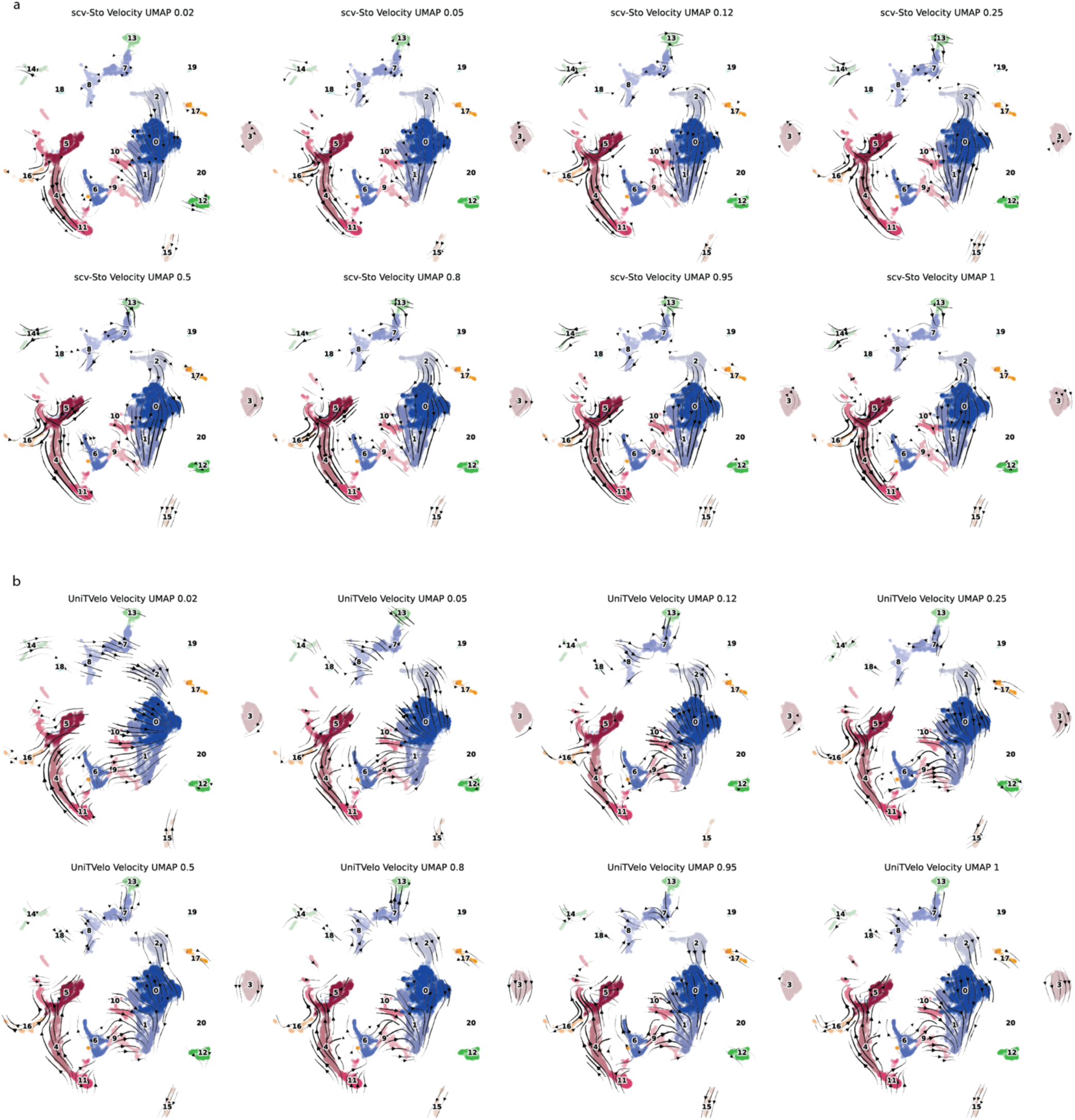
Velocity UMAPs with Subsets of Reads for scv-Sto and UniTVelo, associated with Figure 5. a. UMAP embeddings of the ZF 24hpf whole-embryo with RNA velocity predictions from scv-Sto calculated from subsets 2, 5, 12, 25, 50, 80, 95 and 100% of the reads. b. UMAP embeddings of the ZF 24hpf whole-embryo with RNA velocity predictions from UniTVelo, calculated from subsets 2, 5, 12, 25, 50, 80, 95 and 100% of the reads.

**Supplementary Figure 10.**
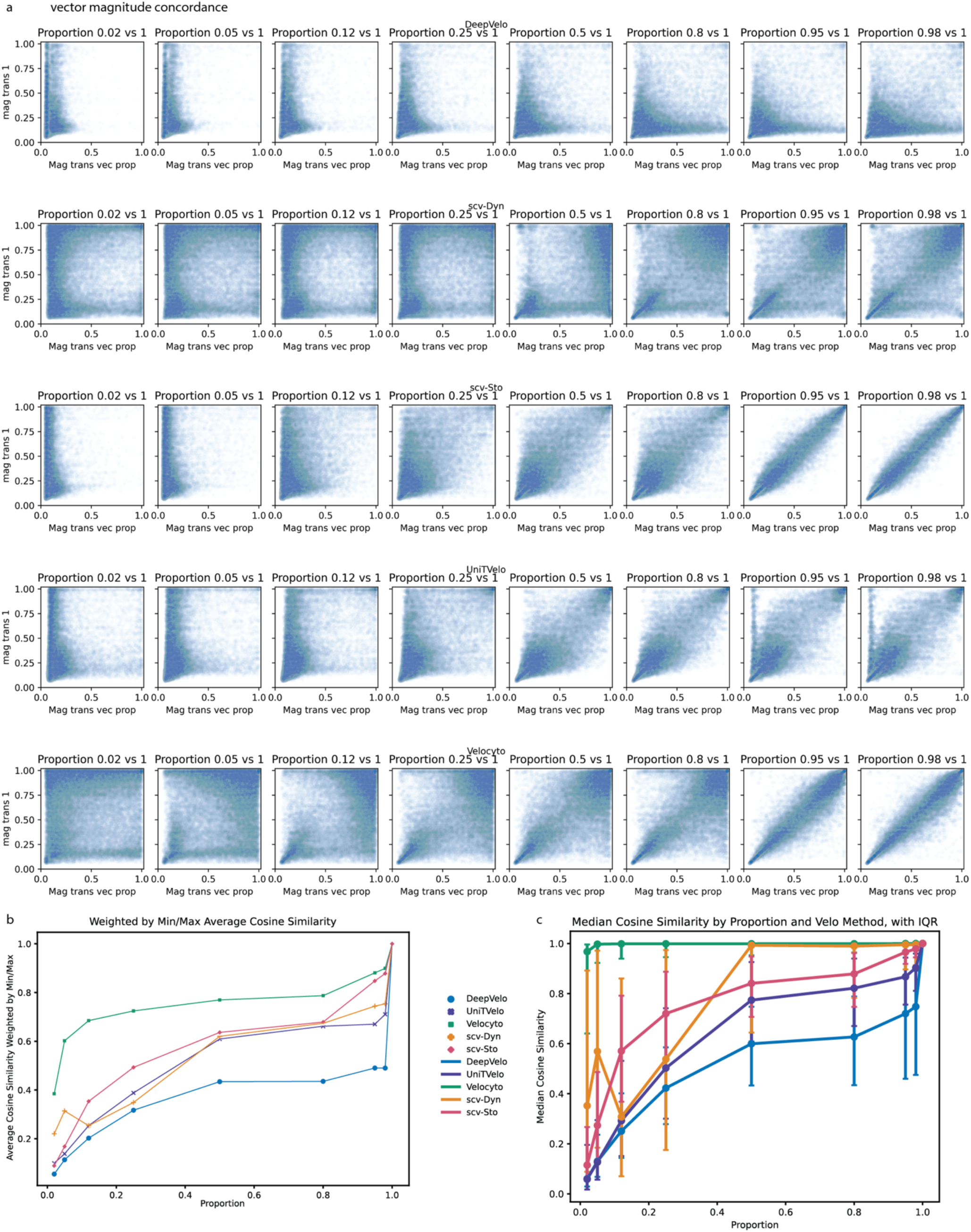
Robustness comparison of magnitude and direction for subset reads, associated with Figure 5. a. Scatterplots comparing the magnitude of the transition vector calculated from the subset reads (x-axis) vs. 100% of the reads (y-axis) for each method. The columns correspond to different subsets, with increasing proportions of reads (2, 5, 12, 25, 50, 80, 95, 98%), and the rows correspond to different methods (DeepVelo, scv-Dyn, scv-Sto, UniTVelo, Velocyto). b. Comparison of directionality robustness for each method weighted by the min/max, calculated as the cosine similarity of the transition vector from the subset with the directionality from the transition vector calculated with 100% of the reads. We multiply each similarity score by the minimum magnitude of the two vectors divided by the maximum magnitude. The line plot shows the averaged value across all cells and subset iterations in the ZF 24hpf whole-embryo dataset. c. Median and InterQuartile Range (IQR −25% and 75% percentile) of the cosine similarity between the transition vector from the subset with the directionality from the transition vector calculated with 100% of the reads, calculated across all cells and subset iterations in the ZF embryo 24hpf dataset.

## References

1. Wagner, D. E. & Klein, A. M. Lineage tracing meets single-cell omics: opportunities and challenges. Nat. Rev. Genet. 21, 410–427 (2020).

2. Street, K. et al. Slingshot: cell lineage and pseudotime inference for single-cell transcriptomics. BMC Genomics 19, 477 (2018).

3. Trapnell, C. et al. The dynamics and regulators of cell fate decisions are revealed by pseudotemporal ordering of single cells. Nat. Biotechnol. 32, 381–386 (2014).

4. Haghverdi, L., Büttner, M., Wolf, F. A., Buettner, F. & Theis, F. J. Diffusion pseudotime robustly reconstructs lineage branching. Nat. Methods 13, 845–848 (2016).

5. Saelens, W., Cannoodt, R., Todorov, H. & Saeys, Y. A comparison of single-cell trajectory inference methods. Nat. Biotechnol. 37, 547–554 (2019).

6. Wyler, E. et al. Single-cell RNA-sequencing of herpes simplex virus 1-infected cells connects NRF2 activation to an antiviral program. Nat. Commun. 10, 4878 (2019).

7. Kimmel, J. C., Yi, N., Roy, M., Hendrickson, D. G. & Kelley, D. R. Differentiation reveals latent features of aging and an energy barrier in murine myogenesis. Cell Rep. 35, (2021).

8. Aissa, A. F. et al. Single-cell transcriptional changes associated with drug tolerance and response to combination therapies in cancer. Nat. Commun. 12, 1628 (2021).

9. Cuevas-Diaz Duran, R., González-Orozco, J. C., Velasco, I. & Wu, J. Q. Single-cell and single-nuclei RNA sequencing as powerful tools to decipher cellular heterogeneity and dysregulation in neurodegenerative diseases. Front. Cell Dev. Biol. 10, (2022).

10. Rossi, G. et al. Capturing Cardiogenesis in Gastruloids. Cell Stem Cell 28, 230–240.e6 (2021).

11. Gorin, G., Fang, M., Chari, T. & Pachter, L. RNA velocity unraveled. PLOS Comput. Biol. 18, e1010492 (2022).

12. Zheng, S. C., Stein-O’Brien, G., Boukas, L., Goff, L. A. & Hansen, K. D. Pumping the brakes on RNA velocity by understanding and interpreting RNA velocity estimates. Genome Biol. 24, 246 (2023).

13. La Manno, G. et al. RNA velocity of single cells. Nature 560, 494–498 (2018).

14. Bergen, V., Lange, M., Peidli, S., Wolf, F. A. & Theis, F. J. Generalizing RNA velocity to transient cell states through dynamical modeling. Nat. Biotechnol. 38, 1408–1414 (2020).

15. Lange, M. et al. CellRank for directed single-cell fate mapping. Nat. Methods 19, 159–170 (2022).

16. Bergen, V., Soldatov, R. A., Kharchenko, P. V. & Theis, F. J. RNA velocity—current challenges and future perspectives. Mol. Syst. Biol. 17, e10282 (2021).

17. Gao, M., Qiao, C. & Huang, Y. UniTVelo: temporally unified RNA velocity reinforces single-cell trajectory inference. Nat. Commun. 13, 6586 (2022).

18. Cui, H. et al. DeepVelo: deep learning extends RNA velocity to multi-lineage systems with cell-specific kinetics. Genome Biol. 25, 27 (2024).

19. Bonner-Weir, S. et al. The pancreatic ductal epithelium serves as a potential pool of progenitor cells. Pediatr. Diabetes 5 Suppl 2, 16–22 (2004).

20. Bastidas-Ponce, A. et al. Comprehensive single cell mRNA profiling reveals a detailed roadmap for pancreatic endocrinogenesis. Development 146, dev173849 (2019).

21. Lange, M. et al. Zebrahub – Multimodal Zebrafish Developmental Atlas Reveals the State-Transition Dynamics of Late-Vertebrate Pluripotent Axial Progenitors. 2023.03.06.531398 Preprint at 10.1101/2023.03.06.531398 (2023).

22. Qiao, C. & Huang, Y. Representation learning of RNA velocity reveals robust cell transitions. Proc. Natl. Acad. Sci. 118, e2105859118 (2021).

23. Li, J., Pan, X., Yuan, Y. & Shen, H.-B. TFvelo: gene regulation inspired RNA velocity estimation. 2023.07.12.548785 Preprint at 10.1101/2023.07.12.548785 (2023).

24. Li, T. On the Mathematics of RNA Velocity I: Theoretical Analysis. *CSIAM Trans*. Appl. Math. 2, 1– 55 (2021).

25. Bayner, J. S. Developmental Biology. Eleventh Edition. Yale J. Biol. Med. 90, 697–698 (2017).

26. Fior, R. et al. The differentiation and movement of presomitic mesoderm progenitor cells are controlled by Mesogenin 1. Dev. Camb. Engl. 139, 4656–4665 (2012).

27. Sambasivan, R. & Steventon, B. Neuromesodermal Progenitors: A Basis for Robust Axial Patterning in Development and Evolution. Front. Cell Dev. Biol. 8, (2021).

28. Ebrahim, N., Shakirova, K. & Dashinimaev, E. PDX1 is the cornerstone of pancreatic **β**-cell functions and identity. Front. Mol. Biosci. 9, 1091757 (2022).

29. Marot-Lassauzaie, V. et al. Towards reliable quantification of cell state velocities. PLOS Comput. Biol. 18, e1010031 (2022).

30. Luecken, M. D. et al. Benchmarking atlas-level data integration in single-cell genomics. Nat. Methods 19, 41–50 (2022).

31. Li, C., Virgilio, M. C., Collins, K. L. & Welch, J. D. Multiomic single-cell velocity models epigenome-transcriptome interactions and improves cell fate prediction. Nat. Biotechnol. 41, 387–398 (2023).

32. Wu, S. & Schmitz, U. Single-cell and long-read sequencing to enhance modelling of splicing and cell-fate determination. Comput. Struct. Biotechnol. J. 21, 2373–2380 (2023).

33. Zhang, C. et al. Improving the RNA velocity approach with single-cell RNA lifecycle (nascent, mature and degrading RNAs) sequencing technologies. Nucleic Acids Res. 51, e112 (2023).

34. Lex, A., Gehlenborg, N., Strobelt, H., Vuillemot, R. & Pfister, H. UpSet: Visualization of Intersecting Sets. IEEE Trans. Vis. Comput. Graph. 20, 1983–1992 (2014).

